# Efficient consideration of coordinated water molecules improves computational protein-protein and protein-ligand docking

**DOI:** 10.1101/618603

**Authors:** Ryan E. Pavlovicz, Hahnbeom Park, Frank DiMaio

**Affiliations:** Department of Biochemistry, University of Washington, Seattle, WA 98195, United States; Institute for Protein Design, University of Washington, Seattle, WA 98195, United States

**Keywords:** energy function, force field, implicit water model, explicit water model, binding free energy

## Abstract

Highly-coordinated water molecules are frequently an integral part of protein-protein and protein-ligand interfaces. We introduce an updated energy model that efficiently captures the energetic effects of these highly-coordinated water molecules on the surfaces of proteins. A two-stage protocol is developed in which polar groups arranged in geometries suitable for water placement are first identified, then a modified Monte Carlo simulation allows highly coordinated waters to be placed on the surface of a protein while simultaneously sampling amino acid side chain orientations. This “semi-explicit” water model is implemented in Rosetta and is suitable for both structure prediction and protein design. We show that our new approach and energy - model yield significant improvements in native structure recovery of protein-protein and protein-ligand docking.

## Introduction

Water plays a significant role in biomolecular structure. The hydrophobic effect drives the collapse of proteins into their general shape while well-coordinated water molecules (water molecules making multiple water-protein hydrogen bonds) on the surface of a protein may confer specific conformations to nearby polar groups. Furthermore, water plays a key role in biomolecular recognition: when a ligand binds its host in an aqueous environment, it must displace water molecules on the surface and energetically compensate for the lost interactions. Coordinated water molecules may also drive host-ligand recognition by bridging interactions between polar groups on each side of the complex.

Simulations of proteins in explicit solvent have been successful in predicting folded conformations^1^ as well as computing binding free energies^2^ with high accuracy. This comes at significant computational cost, while the use of implicit solvent^3^ greatly expedites such calculations, but at the loss of accuracy achieved through the inclusion of highly-coordinated water molecules^4^. Thus, an approach combining the efficiency of implicit solvation with the ability to recapitulate well-coordinated water molecules is desired. Several such methods have been developed but tend to be target-specific^5–8^ or relatively expensive computationally^9–10^.

In this paper, we describe the development of general methods for capturing the energetic effects of explicit solvent, but with the computational efficiency of an implicit solvent model, making the approach suitable for protein-protein and protein-ligand docking. The methods include: 1.) a new energy function that implicitly captures the energetics of protein and coordinated-water interactions and 2.) a conformational sampling approach that efficiently samples protein and explicit water conformations simultaneously. This approach yields superior results in predicting coordinated water positions as well as improving the ability to discriminate native protein-protein and protein-ligand interfaces from decoys.

## Results

Our approach for modeling coordinated water molecules using Rosetta, fully described in *Methods*, is briefly presented here. We have developed two complimentary approaches for capturing coordinated-water energetics. First, *Rosetta-ICO (Implicit Consideration of cOordinated water)*, implicitly captures pairs of polar groups arranged such that a theoretical “bridging” water molecule may form favorable hydrogen bonds to stabilize the interaction. This calculation is efficient but ignores multi-body interactions that may favor, for example, waters coordinated by >2 hydrogen bond donors or acceptors. Therefore, we have also developed *Rosetta-ECO (Explicit Consideration of cOordinated water)*, in which Rosetta’s Monte Carlo (MC) simulation is augmented with moves to add or remove explicit solvent molecules from bulk. By sampling water orientations at sites where predicted bridging waters overlap (Figure 1), we properly coordinate water molecules to optimize hydrogen bonding.

**Figure 1.**
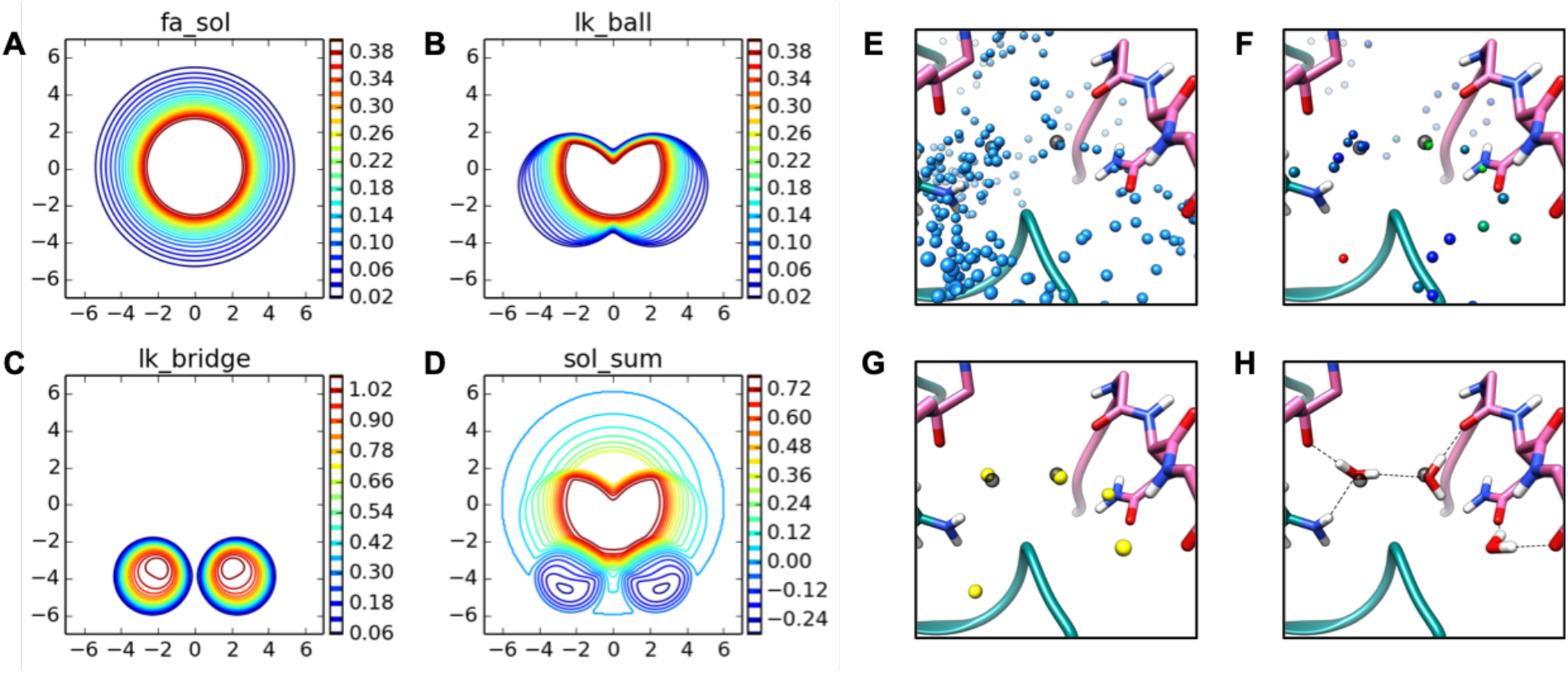
Implicit and Explicit Treatment of Water In Rosetta. Implicit water score function potentials, panels. **A-D.** Potential plots were generated by orienting the N-H and C=O groups of two ALA residues along the same axis with a H--O distance of 1.3 Å (origin). The donor residue is then shifted +/-7 A to generate a planar cut of the solvation potentials between the N and O atoms. **A.** fa_sol term: isotropic desolvation penalty implemented in Rosetta using the Lazaridis-Karplus model. **B.** lk_ball term: anisotropic correction for polar atom types, first introduced into the REF2015 score function. **C.** lk_bridge term: anisotropic solvation reward introduced into the *Rosetta-ICO* score function. **D.** Composite of panels A-C, using the finalized *Rosetta-ICO* score term weights: sol_sum = 1.0*fa_sol + 0.92*lk_ball - 0.33*lk_bridge. Panels A-C show unweighted potentials. **Explicit water placement with *Rosetta-ECO*, Panels E-H. E.** Initial possible solvation sites (blue) are based on statistics of water positions about backbone polar atoms in addition to sites about side chain polar atoms based on all available rotameric positions. Pictured is the interface of PDB ID: 1P57, between the N-terminal (pink) and catalytic (teal) domains of hepsin, with crystallographic waters in transparent grey. **F.** After an initial stage of Monte Carlo packing of both the possible water sites and surrounding protein side chains, a cutoff is applied based on the dwell time of water sites (colored from blue (dwell time = 0%) to green (dwell time = 25%) to red (dwell time = 50%). **G.** Remaining water sites are clustered and a second cumulative dwell time cutoff is applied. **H.** The remaining predicted water sites are converted into three-site water molecules and re-packed with the surrounding side chains using the full Rosetta score function. Two of the final predicted water molecules in this figure are within 0.50 and 0.18 Å of crystallographic water positions, while another water molecule is well-coordinated by the protein, but is not observed in the crystal structure.

For both approaches, the Rosetta energy function has been reoptimized using the *dualOptE* framework described by Park et al.^11^. In this optimization, several metaparameters describing the shape of the *Rosetta-ICO* potential; several terms controlling the strength and shape of protein-water interactions; and ~50 other per-atom polar parameters were optimized to allow for compensating changes to the new energy terms. Energy function parameters for polar groups, including partial atomic charges, were refit using the same training tasks originally used in the parameterization of the *opt-nov15* energy function^11^, now called REF2015^12^. While all of these parameters were optimized for *Rosetta-ICO*, only a subset of water-specific parameters were refit when devloping the explicit water terms for *Rosetta-ECO*. The results in this section are shown with the updated energy functions compared to baseline tests run using the REF2015 energy function^11^.

### Rotamer and Water Recovery at Protein-Protein Interfaces

A set of 153 native protein-protein interfaces from high-resolution X-ray crystal structures was used to test how well the new energy models perform at simultaneously predicting amino acid side chain conformations and coordinated water sites. These tests involved the re-sampling of side chain conformations of interface residues on a fixed backbone in MC simulations, and evaluating resulting predicted side chains against the deposited density maps. In tests involving semi-explicit water molecules (*Rosetta-ECO*), protein and water simultaneously sample conformational space. A baseline rotamer recovery error of 9.73 ± 0.13% (over three runs) was obtained using the *REF2015* energy function for the 7040 flexible side chains of the test set. A marginal improvement is made with *Rosetta-ICO*, reducing error to 9.52 ± 0.04%. Inclusion of explicit water molecules in this test fails to further decrease the overall rotamer recovery error beyond the improvements observed with *Rosetta-ICO*, with a *Rosetta-ECO* error of 9.59 ± 0.15%, while predicting ~19 explicit water molecules per protein-protein interface. For reference, side chain packing tests that keep all benchmark water molecules (perfect recovery and precision) achieves a rotamer side chain recovery error of 8.36 ± 0.04%, while random perturbation of these waters suggest placement tolerance of ~0.8 Å (Fi. S6).

In addition to measuring side chain rotamer recovery at the protein-protein interfaces, we also analyzed the recovery of water positions found in the high-resolution X-ray crystal structures when implementing the *Rosetta-ECO* solvation method. For water recovery tests, modeled water positions are considered “correct” if they are placed within 0.5 Å of the native water or if they are coordinated by the same polar atoms. *Rosetta-ECO* is able to recover 17.1% of native water molecules with a precision of 17.7%. Details of *Rosetta-ECO water* recovery are shown in Table 1. These tables show that our approach is most effective at predicting “buried” waters (28.3% recovery) and highly-coordinated waters (31.2% of triply-coordinated waters). Unsurprisingly, *Rosetta-ECO* is also much more effective at predicted backbone-coordinated waters, correctly predicting 49.4% of backbone-only coordinated waters. An example of correctly predicted water sites is illustrated in Figure 1D.

**Table 1.** Classification of Predicted Native Waters

| Type<sup>1</sup> | Subset Size | *Rosetta-ECO* |  |
| --- | --- | --- | --- |
|  |  | % recovered<sup>2</sup> | % precision<sup>3</sup> |
| All | 3226 | 17.1 | 17.7 |
| Exposed | 773 | 6.0 | 5.0 |
| Partially Buried | 2046 | 19.2 | 22.0 |
| Buried | 407 | 28.3 | 28.3 |
| 1 protein coord | 892 | 6.1 | 6.6 |
| 2 protein coord | 1219 | 26.5 | 26.8 |
| 3 protein coord | 458 | 31.2 | 20.0 |
| BB only | 818 | 49.4 | 10.0 |
| SC only | 814 | 7.0 | 11.3 |
| BB+SC | 1070 | 27.7 | 26.1 |

<sup>1</sup>Three groups of categorization of type of predicted water molecules. First, waters are classified ‘buriedness’ based on number of amino acid neighbors (nC $\beta$ ) with C $\beta$ within 10
Å. Exposed: $nC\beta \leq 15$ ; partially buried: $15 < nC\beta \leq 25$ ; buried: $nC\beta > 25$ . Second, classification by 1, 2, or 3 protein coordination partners within 3.2 Å. Finally, by type of coordinating protein atoms with 3.2 Å of the water O atom: at least two backbone only (BB only), side chain only (SC only) or a mix of backbone and side chain coordination (BB+SC).
<sup>2-3</sup>Percent and number of specific types of waters recovered using recovery criteria described in *Methods*, averaged over three runs.

Finally, the results of *Rosetta-ECO* were compared against solvent placement using the 3D-RISM methodology as implement in AmberTools19^13^. 3D-RISM, like most other water site prediction methods, operates on a fixed structure (the crystallographic structures for this benchmark). In our tests, 3D-RISM recovered 5% more of the full interface water data set than *ECO* when calibrated to the same level of precision (See Table S2 for detailed results). *Rosetta-ECO*, which predicts water positions in addition to protein side chain conformations, performs particularly strongly at recovering waters that are exclusively coordinated by backbone groups (Table 1), outperforming 3D-RISM by 35% for this classification of water. Overall, the 3D-RISM calculations take ~10-fold longer to run.

### Native Interface Recapitulation

We next tested the ability the new energy model to recapitulate near-native conformations of protein-protein interfaces (PPIs) and protein-ligand interfaces. In these tests, the binding free energies for a number of near-native and incorrect (decoy) docking conformations of each complex are computed with the aim of discriminating the correct binding poses from the decoys. PPI decoys were sampled using a combination of Zdock^15^ and RosettaDock^16^, while protein-ligand decoys were generated using RosettaLigand^17^. Both datasets were enriched for water-rich interfaces, leading to 53 protein-protein interfaces and 46 protein-ligand interfaces. Then predicted binding free energies, ΔG_bind_ are calculated for all decoys (see *Methods*). We assess the ability to predict the near-native conformations using a “discrimination score,”^11^ which computes the Boltzmann weight of near-native structures. The values range from 0 to 1, with higher values showing better discrimination. An overview of the results is shown in Table 2.

**Table 2.** Performance of Different Solvation Schemes on Protein-Protein and Protein-Small Molecule Docking Discrimination

|  | REF2105 | Rosetta-ICO<sup>1</sup> | Rosetta-ECO<sup>2</sup> |
| --- | --- | --- | --- |
| *Protein-small molecule* |  |  |  |
| discrimination score<sup>3</sup> | 0.7493 ± 0.0027 | 0.8069 ± 0.0022 | 0.8728 ± 0.0028 |
| percent correct<sup>4</sup> | 77.1 ± 2.1 | 77.8 ± 1.8 | 94.1 ± 1.1 |
| run time<sup>5</sup> | 1.00 | 1.09 | 1.52 |
| Protein-protein |  |  |  |
| discrimination score | 0.6277 ± 0.0138 | 0.7386 ± 0.0061 | 0.7941 ± 0.0038 |
| percent correct | 63.6 ± 0.9 | 74.9 ± 0.9 | 79.9 ± 2.3 |
| normalized run time | 1.00 | 1.25 | 2.59 |

<sup>1</sup>Implicit consideration of coordinated water molecules
<sup>2</sup>Inclusion of well-ordered explicit water molecules
<sup>3</sup>Reported are the average Boltzmann-weighted discrimination scores ± 1σ averaged over three independent runs for 46 protein-ligand and 53 protein-protein docking cases.
<sup>4</sup>The percentage of cases in which the lowest scoring model is within 1.0 Å of the native conformation for protein-ligand docking and 2.0 Å for protein-protein docking, averaged over 3 independent runs
<sup>5</sup>Run time, normalized to baseline, is the sum of individual run times to calculate $\Delta G_{\text{bind}}$ for each near-native and decoy conformation

### Protein-Protein Docking Discrimination

In protein-protein docking discrimination tests, run on a set binding modes that broadly sample RMSD space with respect to the native conformation, significant improvements are observed when comparing *Rosetta-ICO* to the baseline results, with the discrimination score increasing from 0.63 to 0.74. *Rosetta-ECO* further improves this discrimination score to 0.79. We also consider the “success rate,” the time the lowest-energy conformation is within 2.0 Å of native: the *ECO* model enables successful prediction of a near-native conformation in 8 additional cases out of the set of 53, a ~15% improvement. This comes at a modest increase in computational cost, with an average 1.25- and 2.59-fold increase in runtime for *ICO* and ECO, respectively.

As illustrated in Figure 2A, *Rosetta-ECO* improves the discrimination score for 38 of 53 cases, adding 13.4 water molecules to the average bound state and 15.0 water molecules to the average unbound state. These average improvements remain statistically significant. Looking at one such case (adrenodoxin reductase/adrenodoxin, PDB ID 1E6E), we see that while all three energy models correctly predict a near-native conformation, the “energy gap” between native and non-native conformations is improved under *Rosetta-ECO* (Fig. 2B). Closer investigation of the near-native models shows 21 explicit water molecules added to the binding interface. The combined electrostatic and hydrogen bond energy contributions compose a large proportion of the improved binding energy, 5.2 kcal/mol more favorable than *Rosetta-ICO* for this particular binding configuration.

**Figure 2.**
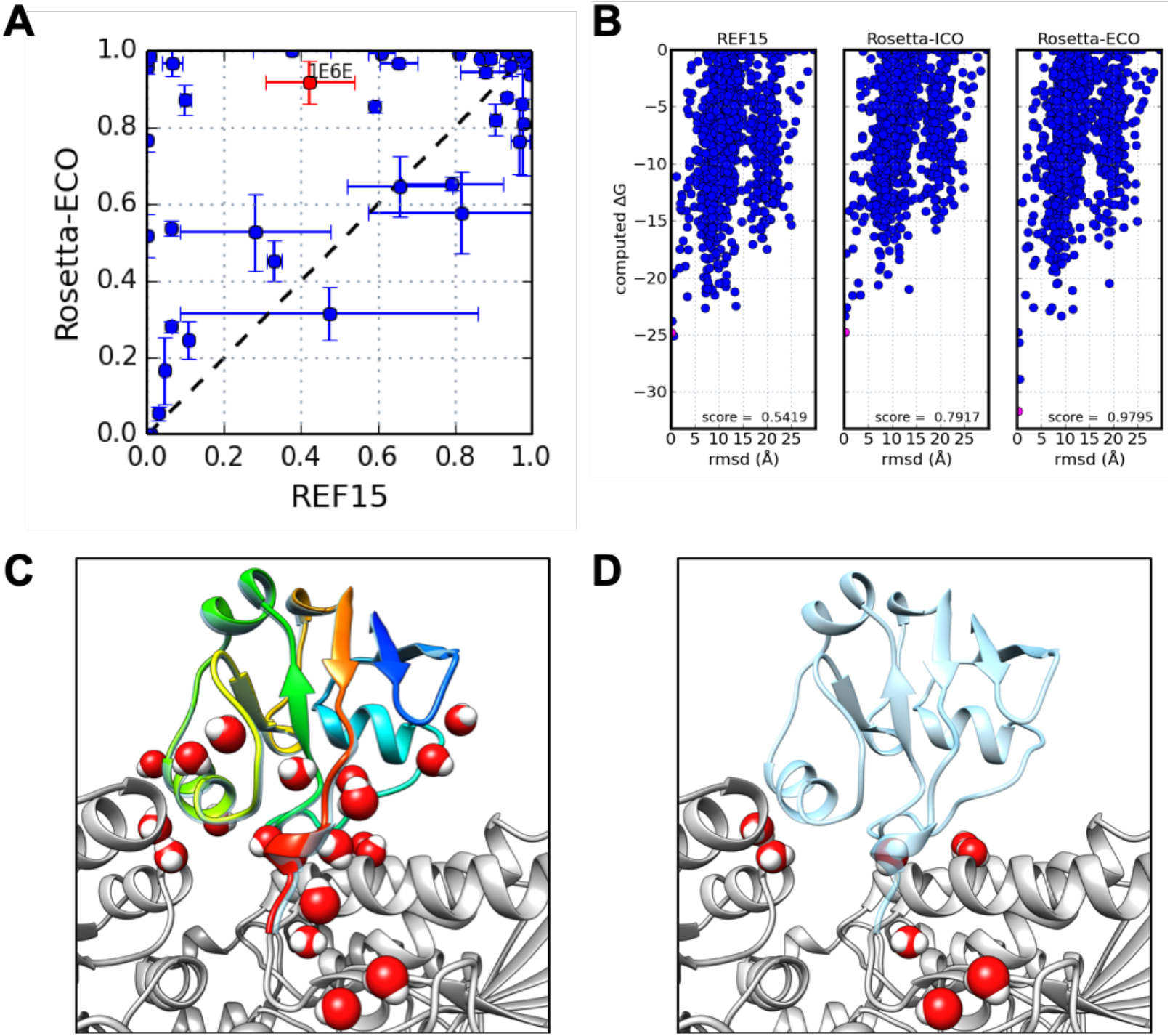
Protein-Protein Docking Results. **A.** Scatter plot comparing results of 53 cases between *REF2015* and *Rosetta-ECO*. Values are the Boltzmann-weighted score ± 1σ from an average of three independent runs. **B.** Energy funnels for PDB ID: 1E6E, adrenodoxin reductase bound to adrenodoxin (red data point in 2A), plotting computed AGbind vs. rmsd from the native binding conformation for three different scoring methods. Boltzmann-weighted scores for each distribution are noted in bottom right of each plot. **C.** Explicitly-solvated near-native docking pose (rmsd=0.14 Å pink data point in 2B) with the reductase in grey and adrenodoxin in rainbow (N-to C-terminus colored blue to red). **D.** The predicted unbound state in which a number of interface waters return to bulk after recalculation. The native ligand binding mode (transparent blue) is shown for reference.

### Protein-Ligand Docking Discrimination

For protein-ligand docking discrimination tests, *Rosetta-ICO* again shows an improvement over *REF2015*, with average discrimination score increasing from 0.75 to 0.81. *Rosetta-ECO* further increases the discrimination score to 0.87. In terms of “success rate”, we see the same trend as with PPIs: *Rosetta-ECO* enables the correct prediction (within 1.0Å of native) in 7 additional cases out of the 46. These results indicate that both *Rosetta-ICO* and *ECO* help discriminate distant decoys from native conformations when compared to the *REF2015* energy model, with the inclusion of explicit water modeling in *ECO* conferring the largest benefit. This also comes at only a modest increase in run time: about 10% increased time for ICO, and about 52% increased computation time for ECO.

The improvements in discrimination score on a case-by-case basis are illustrated in Figure 3A. Here, we see that *Rosetta-ECO* provides a near across-the-board improvement in native discrimination compared to the baseline calculations. The individual energy distributions for PDB ID 1X8X (tyrosyl t-RNA synthase / tyrosine) in Figure 3B show how both *REF2015* and *Rosetta-ICO* incorrectly favor a decoy 6.6 Å from native. *Rosetta-ECO’s* explicit waters dramatically alter the binding energy landscape, improving the discrimination score from 0.27 to 0.89, and energetically favoring a structure only 0.43 Å from native. The *ECO* model predicts two water molecules that bridge the carboxyl group of the tyrosine ligand to interactions with and arginine side chain and a backbone nitrogen group (Fig. 3C): these provide favorable interactions to the native state, with electrostatic and hydrogen bonding interactions a combined 7.5 kcal/mol more favorable when including the explicit interface waters in the *ECO* calculations.

**Figure 3.**
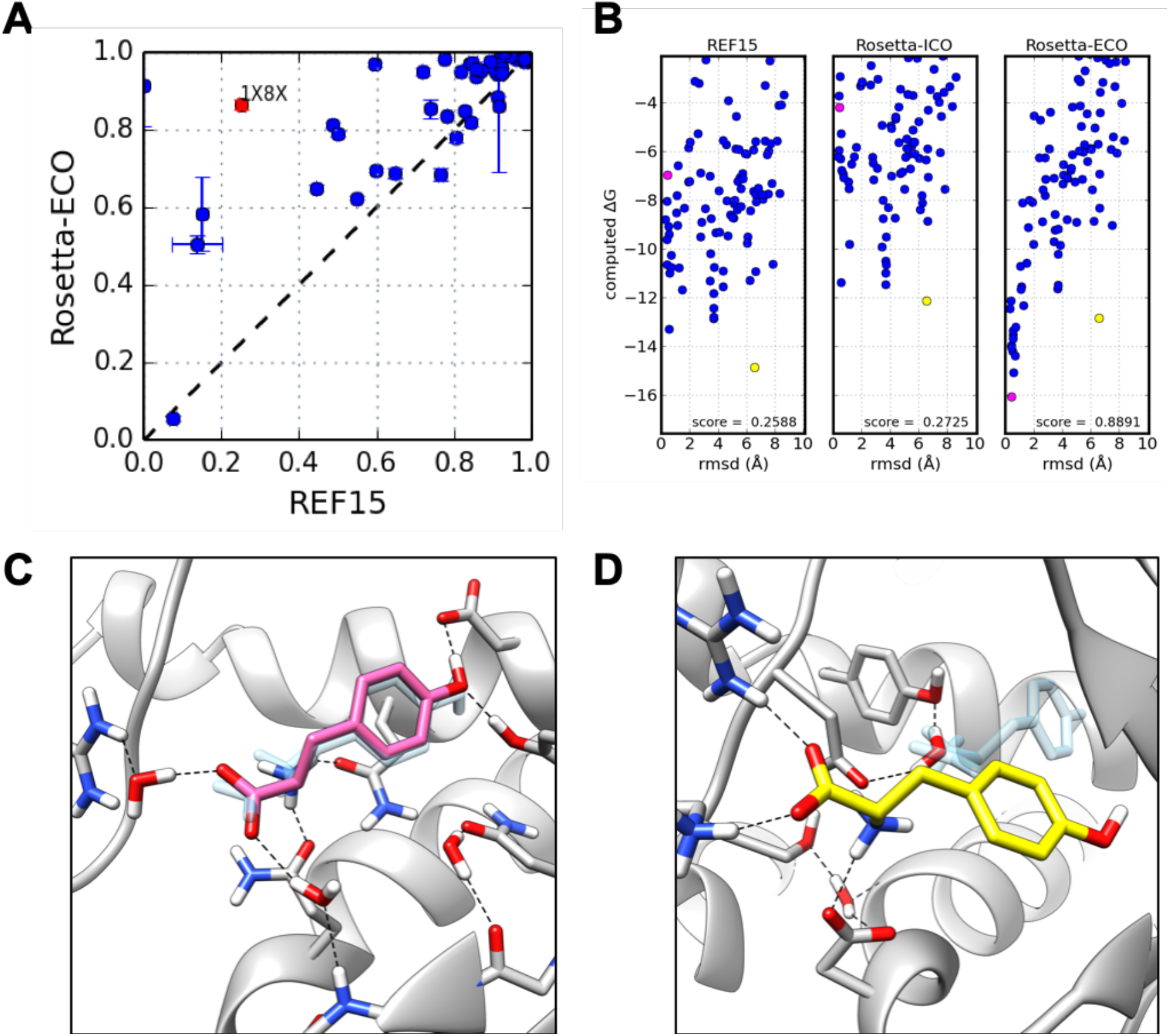
Protein-Ligand Docking Results. **A.** Scatter plot comparing results of 46 cases between baseline *(REF2015)* and *Rosetta-ECO*. Values are the Boltzmann-weighted score ± 1σ from an average of three independent runs. **B.** Energy funnels, similar to Figure 2, for PDB ID: 1X8X, tyrosyl t-RNA synthase bound to tyrosine (red data point in 3A) **C.** Explicitly-solvated, near-native docking pose in pink (RMSD=0.43 Å; pink data point in 3B) with native ligand in transparent blue. **D.** Explicitly-solvated decoy binding pose (RMSD=6.57 Å; yellow data point in 3B).

### Ligand Docking Scoring Comparison

Finally, the new energy functions were compared against the results of a state-of-the art docking approach on a standardized dataset. A recent survey^18^ of widely-used small molecules docking programs tested for performance against the Astex Diverse Set^19^ which includes 85 targets with ligands of pharmaceutical interest. We generated decoys for a 67-target subset, excluding cases in which the ligand is coordinated by an ion, using the top-performing docking software, GOLD^20^. The GOLD-sampled structures were then rescored using the *REF2015*, *ICO*, and *ECO* energy functions of Rosetta. The results, fully presented in Figure S1 and Table S1, show that while the Rosetta-rescored structures are more accurate than GOLD (78.2% versus 67.7% accuracy within a 1 Å RMSD cutoff; 94.6% versus 80.7% accuracy within 2 Å RMSD cutoff), little improvement is observed between *REF2015* and *ICO/ECO*. While these results suggest Rosetta may be a powerful tool for this dataset, the restricted conformational sampling obtained from GOLD (see Figure S2 for examples of sampling in RMSD space) does not benefit from the water model developments presented here and prevents a thorough evaluation of the energy functions. It is likely that a more evenly distributed set of docking conformations would yield results similar to the score function improvements observed in the more tightly-curated protein/protein and protein/ligand data sets described above.

## Discussion

We have presented two approaches for considering coordinated water molecules in the prediction of native protein-protein and protein-ligand interfaces: *Rosetta-ICO*, which very efficiently captures the energetics of bridging waters implicitly, and *Rosetta-ECO*, which allows a small set of waters to emerge from bulk, resulting in a more physically complete representation of protein surfaces and interfaces. Both methods show improvements in protein interface recapitulation tasks with different levels of efficiency/accuracy tradeoffs: *Rosetta-ECO* more accurate but 1.5-2 times slower than *Rosetta-ICO* depending on interface size. The level of native water recovery for *Rosetta-ECO* is about ~5% less than 3D-RISM for a similar precision level, yet the ECO model performs this task at ~10-fold increased speed while simultaneously predicting interface side chain configurations.

Furthermore, while this work highlights the results of water prediction and protein interface recapitulation, we might expect the *Rosetta-ICO* energy function to show modest improvements at tasks related to monomeric structure prediction and protein sequence design. Indeed, that seems to be the case: when tested on independent datasets, modest improvements were observed in decoy discrimination with *ICO*. All other metrics were comparable between the two energy functions, leading us to conclude that the *ICO* model is a reasonable general-purpose energy function.

The improvement in both the protein and ligand docking tests suggests that these new energy functions may prove useful in the design of novel proteins intended to bind a particular ligand or protein. Successful design of protein-protein interfaces is often driven by van der Waals interactions that arise from shape complementarity, however better consideration of ordered solvent molecules may allow for the design of more natural interfaces which include numerous polar residues. Application of these new methods need not be limited to the solvation of interfaces or the description of binding partners. For example, the methods may be applied to more accurately predict the folded state of monomeric proteins in which buried solvent plays an important structural role or for prediction of the stabilizing or destabilizing effect of mutated residues on the surface of a protein. Additionally, the experiments described herein only consider the solvation of proteins and small molecules, however the framework can be easily extended to solvate other biomolecules such as nucleic acids.

## Methods

Two new biomolecular solvation methods are introduced here. The first builds upon the existing implicit water model used in Rosetta to not only account for desolvation penalties, but energetically reward conformations that are suited to accommodate theoretic bridging waters which are calculated on the fly. The second model places well-coordinated water molecules on the surface or at interfaces of biomolecules based largely on statistics from high-resolution experimental data.

### 1.) Implicit Solvation (*Rosetta-ICO*)

An additional energy term is added to the Rosetta’s implicit solvation model that models the energetic costs of highly ordered water molecules coordinated by multiple protein polar groups. The term builds upon our previously developed anisotropic solvation model^11^, where for each polar group, one or more virtual water sites are placed in a configuration ideal for hydrogen bonding with the corresponding polar group. An energetic bonus is then given when the water sites of multiple polar groups overlap in such a way that a single water could coordinate, or “bridge”, these polar groups:

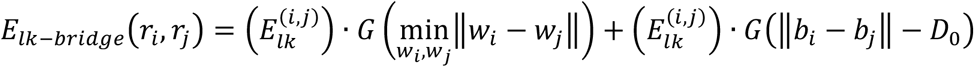

With:

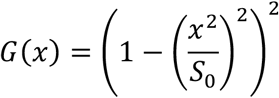

Here, *w_i_* is the xyz coordinate of a theoretic water corresponding to polar group *r_i_*; *b_i_* is the xyz coordinate of the base heavy atom used to construct the water (e.g., the backbone N or O), and D_0_ and S_0_ are parameters that are optimized during energy function evaluation. The two terms in the equation characterize the overlap and the angle formed between polar groups that potentially coordinate a water.

This energy term was added to the current anisotropic solvation model in Rosetta (illustrated in Fig. 1 A-D), and optimization of all polar terms was carried out. While this term does not prevent disallowed coordination geometries (e.g., 3 donors or 3 acceptors coordinating a single water site), in practice, the water sites implicitly identified by this approach are quite reasonable. Because this two-body energy term is only dependent upon the configuration of pairs of protein polar groups, it can be used in all Monte Carlo minimization methods used in Rosetta^21^, with negligible computational overhead.

Additionally, to properly handle the geometry of water-protein and water-water hydrogen bonds, we modified the functional form of sp3-hybridized hydrogen bond acceptors. Previously, the interaction between a hydrogen bond donor and the lone pair electrons of sp3-hybridized acceptors was described by an angle and torsional term about the base atoms; e.g., for serine, the angle CB-OG…Hdon and the pseudo-torsion HG-CB-OG…Hdon. For water, however, this led to an undesirable property in that the potential treated water asymmetrically. Therefore, the torsional term water replaced with a “softmax” potential between the both atoms bonded to the sp3-hybridized acceptor:

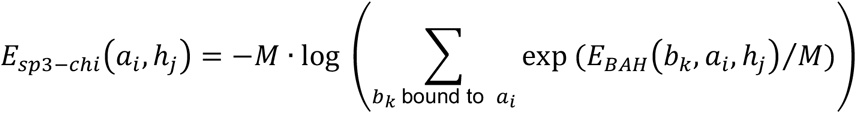

Above, a_i_ and h_j_ are the acceptor heavy-atom and donor hydrogen, respectively; E_BAH_ is the angular potential about the heavy-atom^22^. The summation is carried out over all bound atoms to the acceptor: for water acceptors, this would be over both hydrogens. In the serine example above, the angular potential is applied to both CB-OG- - - Hdon and HG-OG- - - Hdon and the softmax gives a score equal to the worse of the two angular potentials. This ensures the potential is symmetric about both water hydrogens.

### 2.) Explicit Solvation Model (*Rosetta ECO*)

One key challenge in prior explicit water modelling^23^ is the large conformational space a single water molecule can adopt. This is a particular issue in applications (like those in this manuscript) where it is desirable to simultaneously sample side chain conformations and water positions. *Rosetta-ECO* makes use of a two-stage approach to get around this problem (Figure 1). In the first stage, rotationally independent “point waters” are sampled using a statistical potential; not considering water rotation lets thousands of putative water positions be sampled efficiently. In the second stage, for the most favorable water positions (typically only several dozen) we consider rotations of these molecules using a physically derived potential.

In both steps of the protocol, Monte Carlo sampling is used to simultaneously sample side chain and water conformational states. In both stages, water molecules may be set to “bulk,” losing an entropic penalty by doing so. This entropy bonus value, Eref, ultimately controls the number of explicit water molecules placed by the algorithm, requiring sufficient favorable physical interactions to overcome the entropic cost of coming out of bulk. Rotational sampling of waters uses a uniform SO3 gridding strategy^24^ with 30° angular spacing.

#### 2.1) Derivation of the Statistical Point Water Potential

The first step in determining possible water sites involves a low-resolution, statistical water potential to quickly evaluate the interaction between possible water sites and nearby polar groups of biomolecules. This potential, which we are calling the “point water potential”, treats water molecules as simple, uncharged, points with attractive and repulsive Lennard-Jones terms.

The point water potential takes the form of:

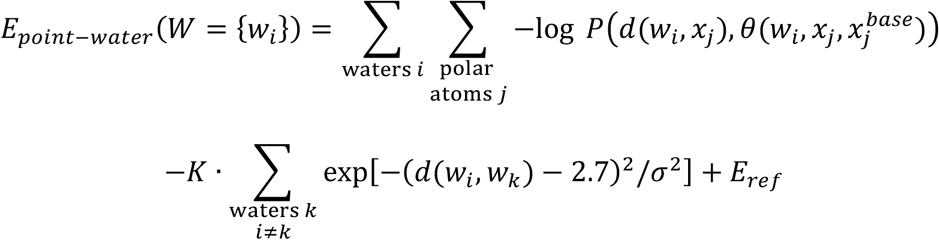

Here, *P* is the statistical point-water distribution, parameterized over distance and angle; *d* gives the distance between a water and polar atom, and *θ* gives the angle between water, polar atom, and its “base atom.” The point water energy term also considers other nearby point water sites, k, as Gaussian distributions with width *σ* and height *K* (with min energy at a distance of 2.7 Å), which was determined by averaging water-water distances observed in high resolution crystal structures. Finally, an overall energetic cost of bringing the water molecule “out of bulk,” E_ref_, is added for each water. These parameters were fit using crystallographic waters in the Top8000 database (see Supplemental for more details).

#### 2.2) Identifying and Packing Point Waters

A key challenging in building possible water sites is we want to simultaneously sample side chain formations along with water positions. Thus, the initial placement of water molecules to be optimized by the point water potential come from two sources: a) ideal solvation about protein backbones, and b) *possible* solvation sites from side chain rotamers. For backbone waters, point generation is straightforward: 10 “ideal” sites are generated from each backbone C=O group (based on clustering waters from crystal structures).

Generation of side chain-coordinated waters is not so straightforward. Considering all possible polar groups of all side chain rotamers is computationally intractable. We again build off prior work^25^ and consider instead side chain/side chain (and side chain/backbone) “overlaps.” That is, we generate all possible side chain rotamers for every side chain, and identify all positions where there is overlap (within 0.75 Å) between two different side chains. A 3D hash table makes this calculation efficient even when there are millions of putative water positions. Finally, to further reduce conformational sampling, during the Monte Carlo “packing” algorithm, when both side chain and point water positions are sampled, all putative point waters are clustered into sets into which only one site can be occupied.

A modified version of Rosetta’s traditional packing algorithm^26^ is used when point waters are present. Typically, Rosetta uses simulated annealing to find the discrete rotamer set minimizing system energy, where the temperature of the trajectory is slowly annealed from RT=100 to RT=0.3. With the point water potential, we do not expect the forcefield (which does not consider water rotation) to be perfect, and we want the packer not to optimize total energy but to simply separate reasonable from unreasonable water positions for a more expensive subsequent calculation. Thus, we instead used long simulations at low temperatures (RT=0.3) with intervening high-temperature “spikes” (RT=100). Then, instead of taking the lowest energy state sampled, we measured water “occupancy” at each position, taking point water positions with a “dwell time” more than 2%.

The water positions passing this criterion, typically only several dozen to a hundred, are then allowed to rotate and are packed (along with all side chains) using Rosetta’s standard simulated annealing rotamer optimization routine.

### 3.) Datasets

Four different data sets were used in the testing of the new energy functions described here. The first includes 153 high-resolution crystal structures of protein-protein interfaces (PPIs) that was used for both native water and rotamer recovery at the interfaces. Two docking data sets were used to test the ability of the new energy functions to discriminate near-native from decoy docking conformations, a subsets of those used by Park et al.^11^, but selected for water-rich interfaces (and to exclude problematic cases such as PPIs with disulfides across the interface or ions contributing to binding). For protein-protein interactions, a 53-case subset of the ZDock 4 Benchmark set^27^ was used, while a 46-case subset of the Binding MOAD database^28^ was used for protein-ligand interactions. Finally, another ligand docking set, generated with GOLD on a subset of the Astex Diverse Set^19^ was used to compare the new energy functions against an established docking score function. Details on the datasets, including lists of PDB IDs used are included in the Supplemental Materials.

### 4.) Benchmarking Against 3D-RISM Water Site Predictions

The water site predictions in Rosetta were compared against those predicted by the 3D-RISM method^29^ as implemented in AmberTools19^13, 30^. Briefly, RISM calculations were performed for pure water at a concentration of 55.5 M with a 0.5 Å grid spacing. Using a buffer of 7 Å, as opposed to the default 14 Å, was found to be speed up calculations while not hurting recovery for our dataset which consists of water molecules found at PPIs. The placevent algorithm^14^ was used to determine explicit water sites, which were truncated to be found within 6 Å of all CB atoms (CA for GLY) of the residues that form the interfaces of the test set. This was done to be comparable to the *Rosetta-ECO* results, which were limited to the PPIs. Finally, the results were further trimmed by the 3D-RISM water-protein radial distribution function (RDF >= 10.2) to achieve the same level of precision as *Rosetta-ECO*.

### 5.) Binding Energy Calculations

The binding free energies, ΔG_bind_, were calculated for the near-native and incorrect (decoy) docking poses by taking the difference between the computed energies of the bound and unbound states. This is accomplished in Rosetta by first calculating the energy for the bound system, then re-computing the energy when the two binding components are separated to obtain unbound state energies. An important part of interface energetics involves computing the energy cost of water displacement^31^, making treatment of explicit waters of the unbound state an important consideration. Due to size differences of the average interface, we found slightly different treatment performed better with PPIs versus protein-ligand interfaces. In both PPIs and protein-ligand interfaces, the bound states are solvated, using the two-stage Monte Carlo procedure described above. This mode of solvation samples both side chain and water orientations, having the effect of considering the induced fit effect. Then all side chains are minimized and, for protein-ligand interfaces only with the *ICO* model, the rigid-body transformation between receptor and ligand is also minimized. Interface components are then separated and re-solvated. Copies of the waters from the bound state are duplicated such that one copy belongs to both ligand and receptor. During the resampling of the unbound state, this allows waters that were previously highly coordinated in the bound state to be liberated to bulk if a sufficient part of this coordination was lost in the unbinding process. Any water molecules that remain un-liberated to bulk following sampling are considered part of the bound/unbound states for scoring purposes.

### 6.) Training Tasks

The training tasks used for energy function parameterization are the same as detailed in the development of the REF2015 Rosetta energy function^11^ and are summarized in the Supplemental Materials.

## ASSOCIATED CONTENT

(Word Style “TE_Supporting_Information”). **Supporting Information**. Additional details on methodology as well as lists of data sets are included in the Supporting Information.

## Author Contributions

The manuscript was written by RP and FD and edited by RP, FD, and HP. RP and FD designed the research, and RP and HP performed research and analyzed results. All authors have given approval to the final version of the manuscript.

## ACKNOWLEDGMENT

This work was facilitated though the use of advanced computational, storage, and networking infrastructure provided by the Hyak supercomputer system at the University of Washington. Structure visualization and analysis used the UCSF Chimera software^32^, while GNU Parallel was used for distributed processing and data analysis^33^.

## ABBREVIATIONS

Rosetta-ICO: implicit consideration of coordinated water
Rosetta-ECO: explicit consideration of coordinated water
PPI: protein-protein interface
RMSD: root-mean-square deviation

## Supporting Data Sets

Data Set 1. Exact PDB files used for water recovery tests.

## Supporting Information

### Dataset Details

#### 1.) Water and Interface Recovery Datasets

A set of 153 high-resolution X-ray crystal structures of protein-protein interactions were used for water prediction and interface rotamer recovery testing. Interface waters were filtered such that each water comes within 3.5 Å of a heavy atom of both chains forming the interface or forms a three-water bridge between the two chains with the anchoring waters coming within 3.5 Å of each chain. An additional filter was used to remove water molecules that clash with one another. Waters within 0.85*2*O_vdw_ of each other, where Ovdw is the van der Waals radius of the water oxygen atom (1.55 Å), were checked for how well they fit into the electron density map of the crystal structure. Any clashing water with an electron density correlation < 3.5 was removed from the final set of interface waters.

Interface residues were defined by the RestrictToInterfaceVector task operation in Rosetta, using the following parameters: vector_dist_cutoff = 9.0; vector_angle_cutoff = 75.0; nearby_atom_cutoff = 5.5; CB_dist_cutoff = 10.0. These residues were solvated with the two-stage method described above. Following the final repack with the full-atom energy function in Rosetta on a fixed backbone, the final computationally-determined side-chain conformations were compared to the experimental structure to measure recovery of rotameric states. Only residues with experimental side-chain conformations that correlate well to the electron density map (correlation score ≥ 0.72) were used for analysis. A total of 7040 side chain rotamers from the test set met this criterion. Predicted side-chain conformations were determined to the native conformation if the difference in electron density correlation was less than or equal to 0.12 with an individual density correlation greater than or equal to 0.71.

Predicted water positions were determined to recall one of the 3290 native interface water molecules if the oxygen position was within 0.5 Å of crystallographic position or if the predicted water coordinates that same neighboring polar groups as a native using a 3.5 Å heavy atom cutoff.

#### 2.) Docking Discrimination Datasets

Two docking data sets were generated for binding energy calculation testing with various score functions: 1.) protein/ligand and 2.) protein/protein docking sets. For both, the goal was to generate ensembles of near-native and decoy binding conformations that were well-distributed in RMSD-space with respect to the experimental conformation. This was achieved through self-docking protocols in addition to perturbation of the native conformation in cases where near-native sampling via docking was poor.

For protein-ligand docking, 46 members of the Binding MOAD database^28^ (those excluding ions or cofactors in the binding sites and enriched for cases in which water molecules are found at the interfaces) were used to evaluate improvements in differentiating native versus decoy docking poses (see Supporting Information for complete list). Ligands parameterized with AM1-BCC partial atomic charges^29^ and locally docked to the native binding site with RosettaLigand^18^. For each ligand, 3000 decoys were generated by docking 30 alternative ligand conformations 100 times. Near-native conformations were generated using the “minimize” option in RosettaLigand^18^ starting with the experimental structure. Finally, after generating an initial docking conformations with the default RosettaLigand energy function, the lower edge of the ΔG_bind_ vs rmsd to native energy distribution was selected as the final test set. This created a total of 6376 total docking conformations including both near-natives and decoys.

For protein-protein docking, 53 cases with a total of 59,738 docking conformations were generated with ZDock 3.0^27^, followed by local optimization with RosettaDock^17^, implementing the same protocol as in *dualOptE*^14^. For both protein-protein and protein-ligand datasets, there is continuous sampling in the RMSD dimension with respect to the native structure.

#### 3.) GOLD Docking Dataset

Ligand docking was carried out to the protein.mol2 files from the Astex Diverse Set available for download from the CCDC website (https://www.ccdc.cam.ac.uk/support-and-resources/downloads/). Prior to docking, ligands and cofactors were assigned partial atomic charges using the AM1-BCC method in *Antechamber^30^*. Each ligand was then docked as described by Liebeschuetz, et al., using the ChemPLP score function with the standard genetic algorithm settings with default early termination parameters which halts the GA when the top three ligand poses are within 1.5 Å of each other^31^. To expand the number of decoys generated by GOLD, the starting point for GOLD docking was randomly perturbed by up to 6.0 Å from the center of geometry of the native ligand position, while maintaining the standard 6.0 Å search radius. The final set used for rescoring with Rosetta includes ~500 docking conformations for a 67 target subset of the Astex Diverse set, excluding the cases in which an ion coordinated the ligand in the binding pocket.

**Figure S1.**
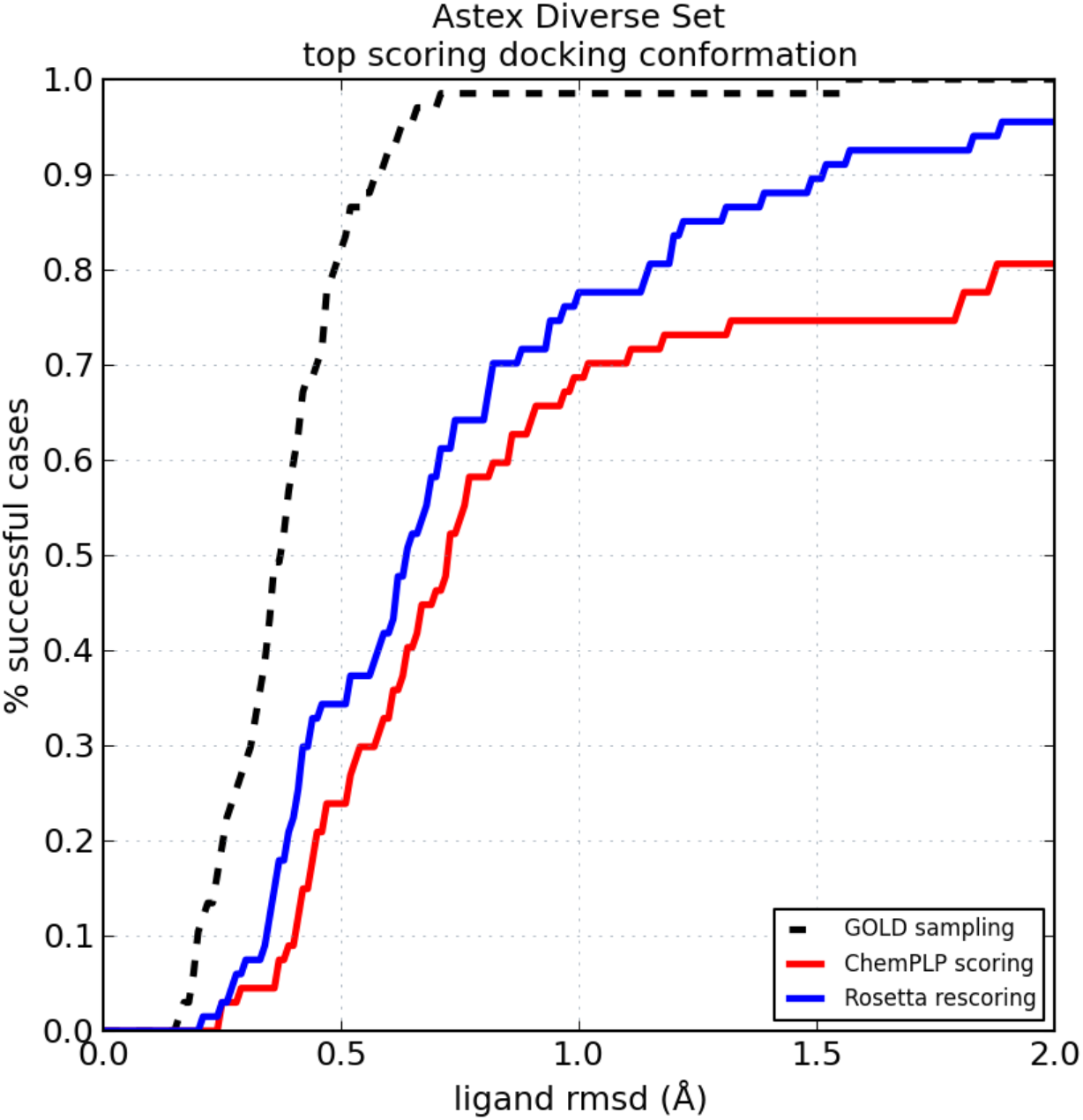
Rescoring GOLD Docking Results With Rosetta. Results for rescoring Astex Diverse Set. Docking conformations initially generated and scored by GOLD (red) were rescored with the Rosetta *REF2015* energy function (blue). The theoretical scoring success is determined by the initial GOLD sampling (black dashed) for the 67 cases of the Astex Diverse Set that do not coordinate an ion in the binding site.

**Table S1.**
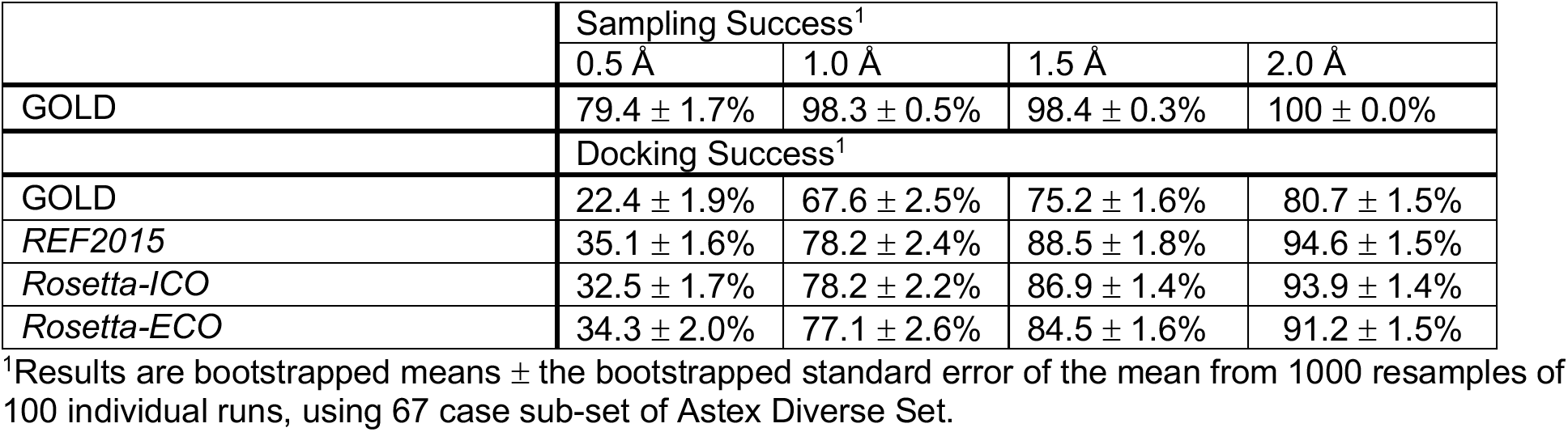
GOLD Docking and Rosetta Rescoring Results of Astex Diverse Set

|  | Sampling Success<sup>1</sup> |  |  |  |
| --- | --- | --- | --- | --- |
|  | 0.5 Å | 1.0 Å | 1.5 Å | 2.0 Å |
| GOLD | 79.4 ± 1.7% | 98.3 ± 0.5% | 98.4 ± 0.3% | 100 ± 0.0% |
|  | Docking Success<sup>1</sup> |  |  |  |
| GOLD | 22.4 ± 1.9% | 67.6 ± 2.5% | 75.2 ± 1.6% | 80.7 ± 1.5% |
| REF2015 | 35.1 ± 1.6% | 78.2 ± 2.4% | 88.5 ± 1.8% | 94.6 ± 1.5% |
| Rosetta-ICO | 32.5 ± 1.7% | 78.2 ± 2.2% | 86.9 ± 1.4% | 93.9 ± 1.4% |
| Rosetta-ECO | 34.3 ± 2.0% | 77.1 ± 2.6% | 84.5 ± 1.6% | 91.2 ± 1.5% |

<sup>1</sup>Results are bootstrapped means ± the bootstrapped standard error of the mean from 1000 resamples of 100 individual runs, using 67 case sub-set of Astex Diverse Set.

**Figure S2.**
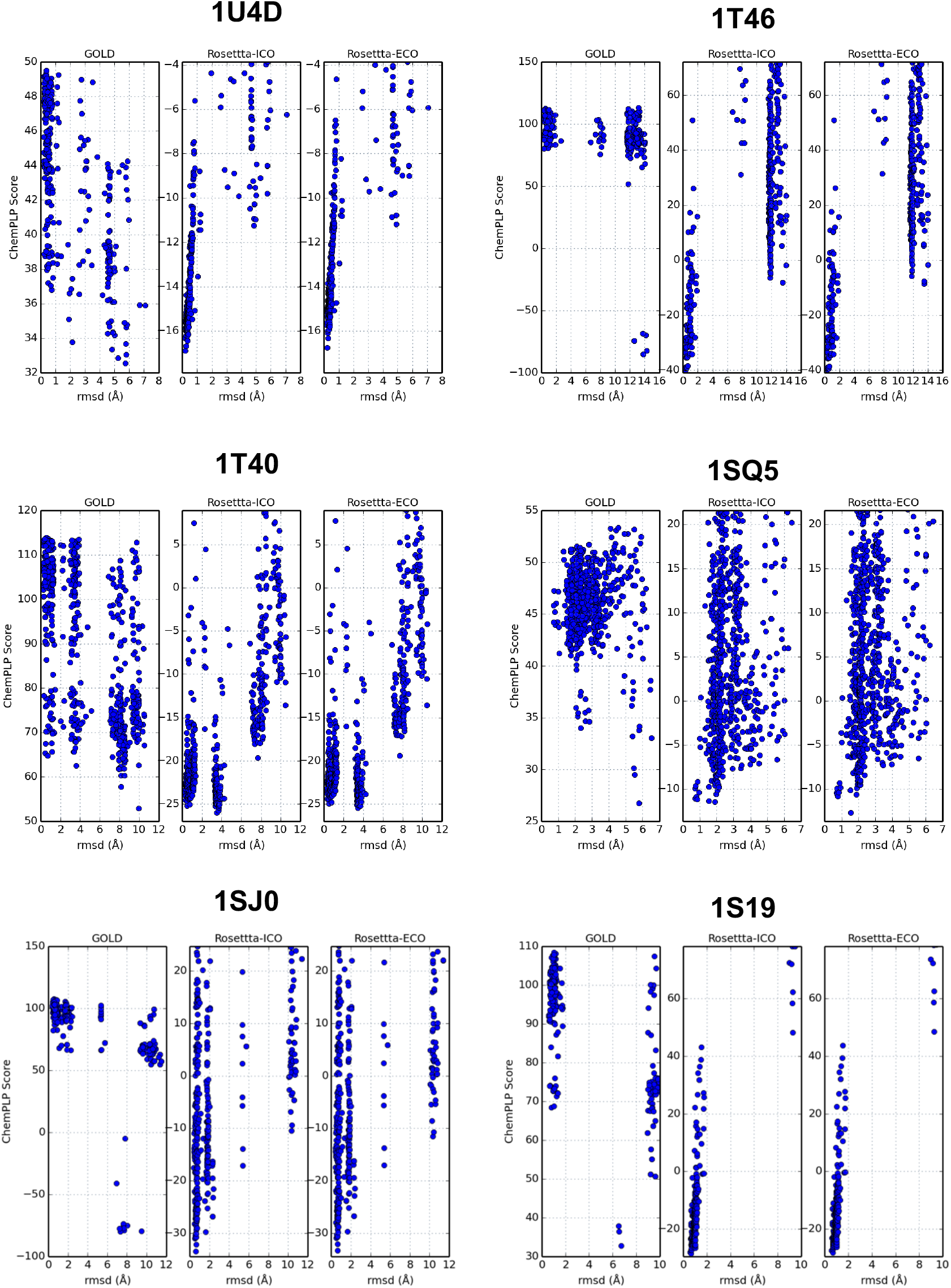
Comparison of Docking Scores/Energies for Conformations Sampled By GOLD for Select Cases. The rmsd of the ligand from the experimental conformation is plotted against the computed score (ChemPLP) for GOLD and ΔG_bind_ for Rosetta. Note that the sampling from GOLD is often focused in small number of docking conformations, leaving gaps in the sampled space.

**Figure S3.**
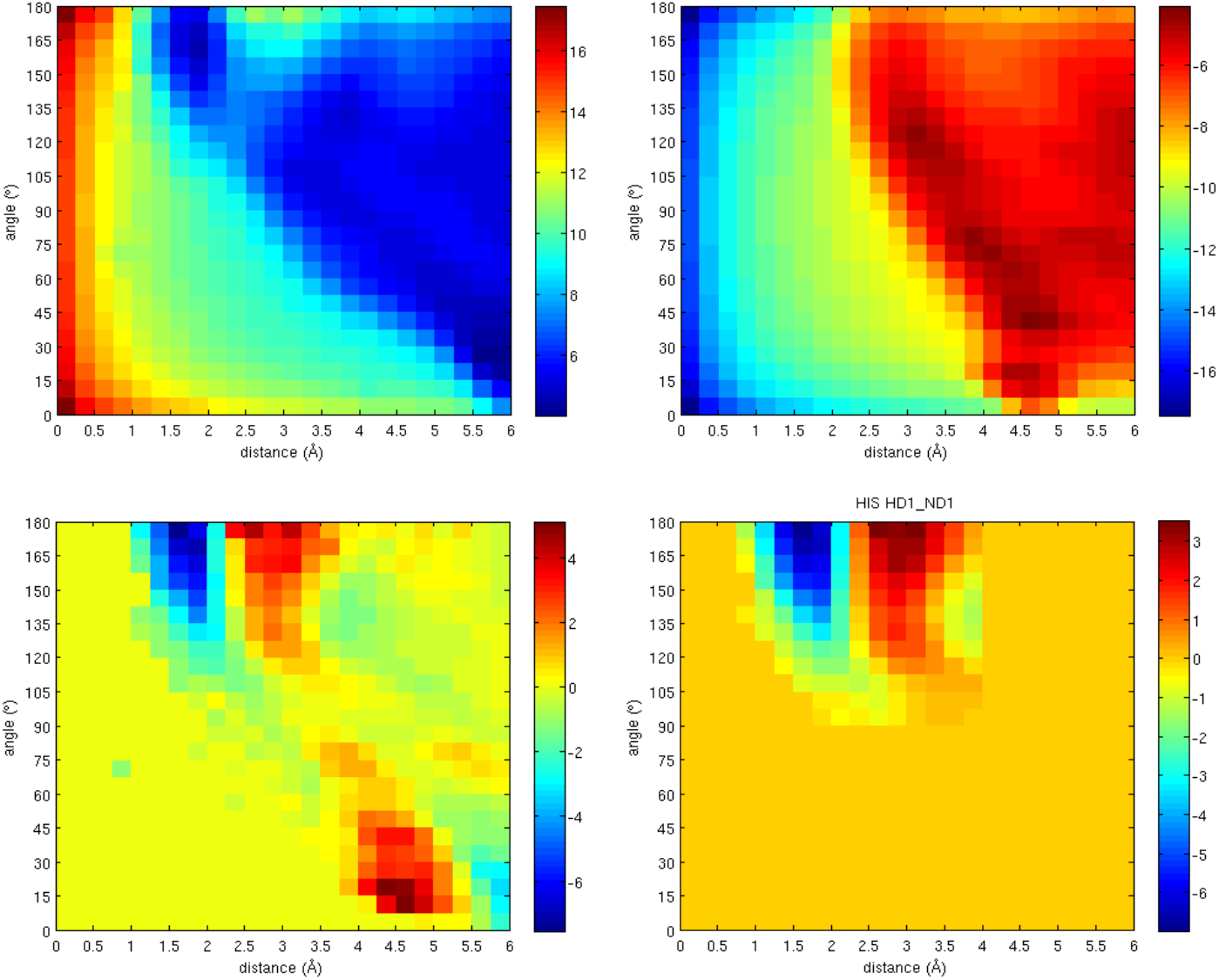
Derivation of a statistical water potential. **Upper left:** Distribution of waters about histidine residues over a range of distance from the HD1 atom and a range of angles from the HD1 and ND1 atoms [-log(HIS_HD1_ND1_)] **Upper right:** Distribution of waters about a non-polar reference [log(ALA_HB1_CB1_)] **Lower left:** The sum of the upper two figures: the statistical potential for histidine **Lower right:** Final, modified histidine potential filtered for noise and second solvation shell effects.

**Figure S4.**
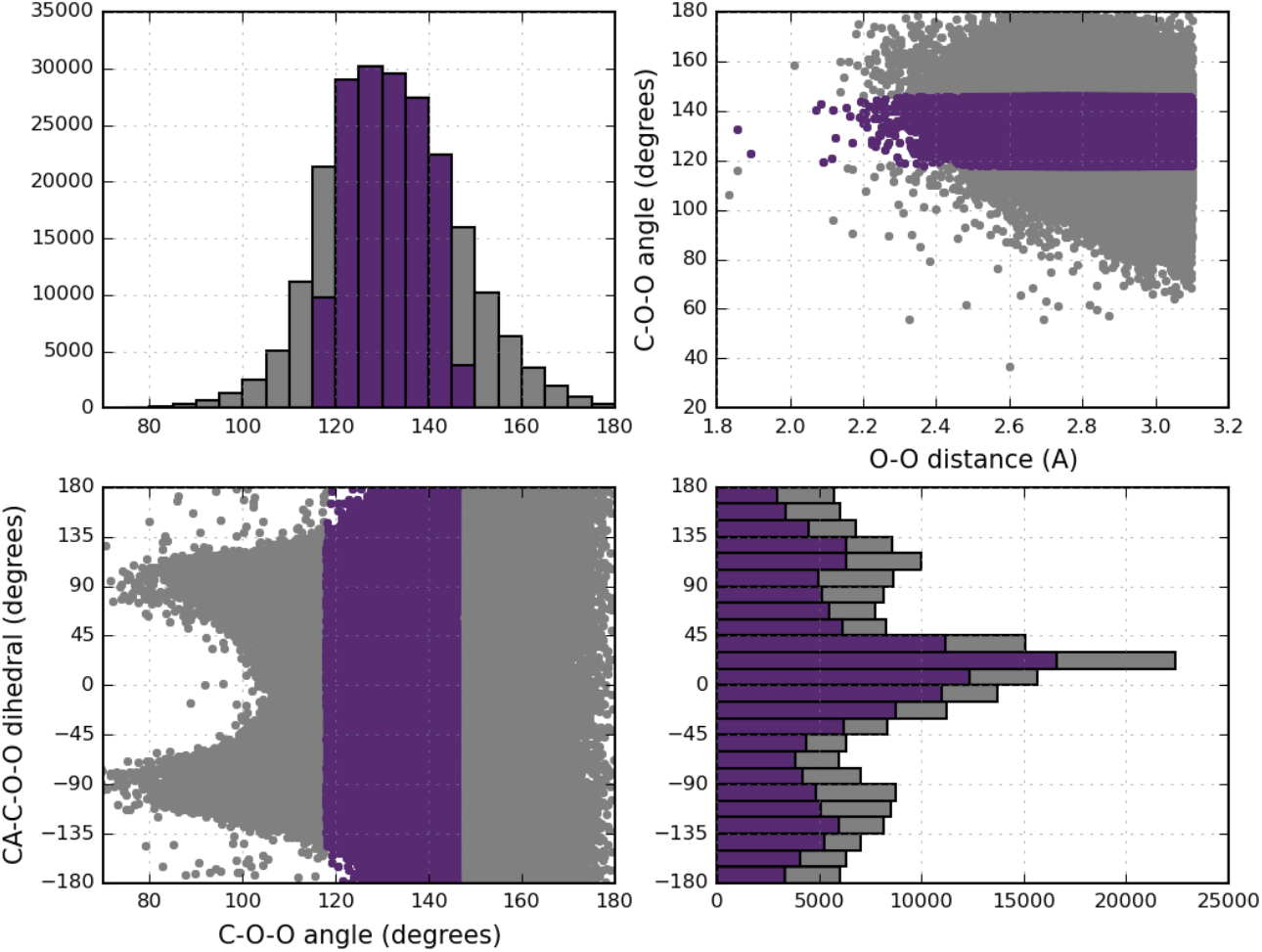
Sample statistics of waters about peptide C=O groups. **Upper right:** distance and angle of all waters measured (grey) and those used for statistical placement about the polar group (purple). **Bottom left:** Angle and dihedral distribution with histogram projections in upper left (angle) and lower right (dihedral).

**Figure S5.**
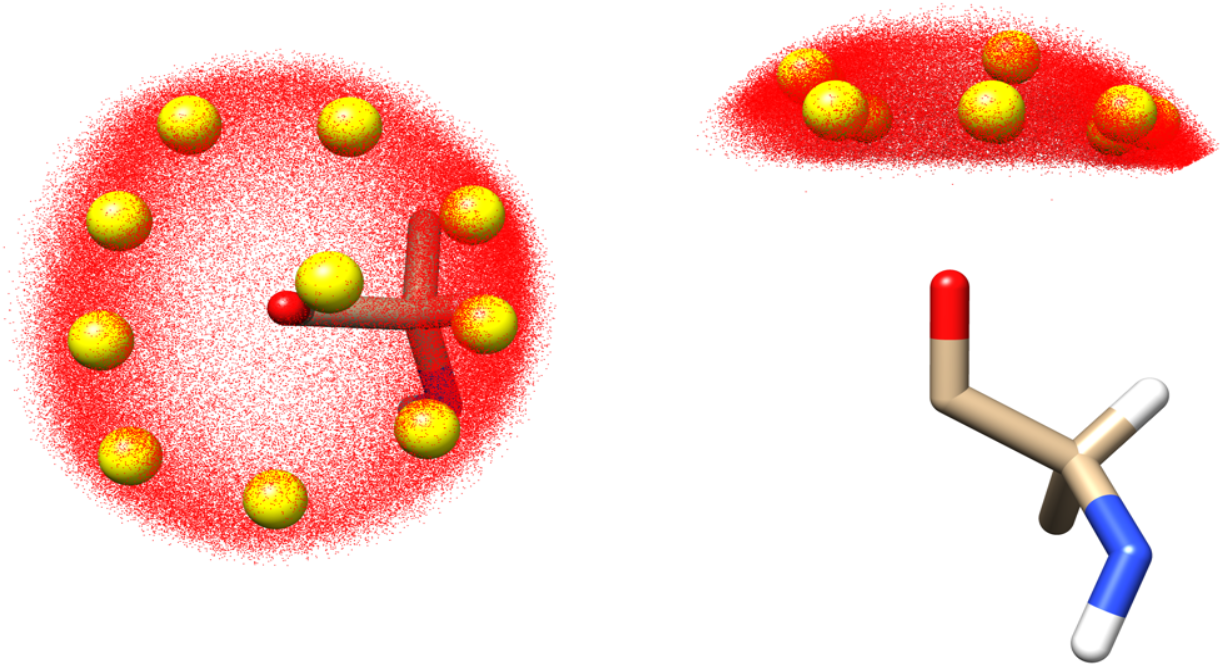
Position of cluster representatives for solvation of C=O backbone groups. The crystallographic water positions used for statistical placement of potential solvation sites about C=O backbone polar groups are shown here in red, with the k-means cluster centroids (k=10) illustrated in yellow. Two views of these data are shown about an arbitrary alanine residue.

**Figure S6.**
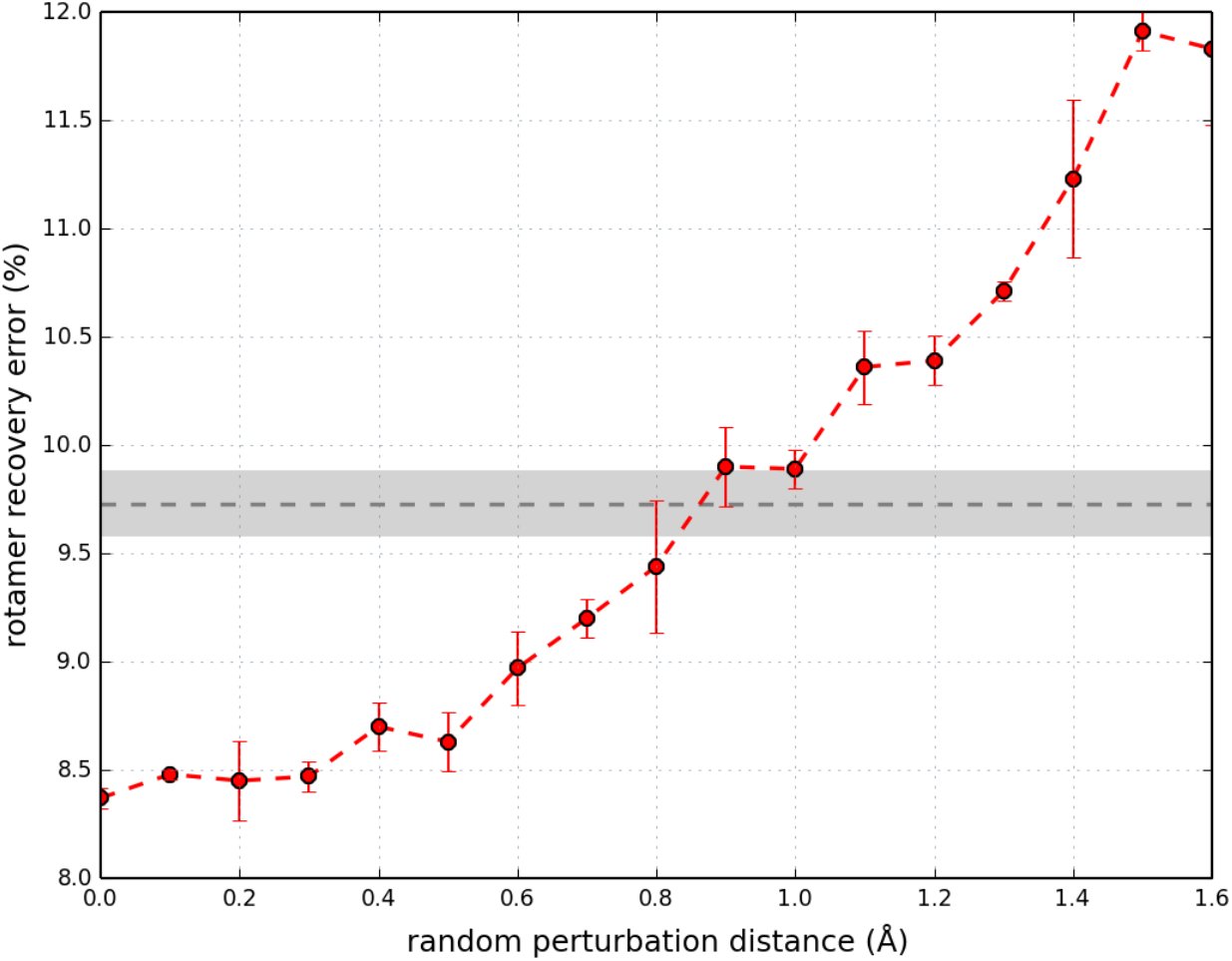
Rotamer recovery error as a function of native water positions randomly perturbed. Crystallographic water molecules in our benchmark set were randomly perturbed 0 to 1.6 Å and the interface residues were repacked in Rosetta. Data points represent the average of three independent runs with 95% confidence interval error bars. The baseline of packing the interfaces without any water molecules (REF2015 score function) is shown as a dashed grey line with 95% confidence intervals from three runs shaded in light grey.

**Table S2.** 3D-RISM Results on Interface Water Test Set.

| Type<sup>2</sup> | Subset Size | 3D-RISM<sup>1</sup> |  |
| --- | --- | --- | --- |
|  |  | % recovered<sup>3</sup> | % precision<sup>4</sup> |
| All | 3226 | 22.9 | 17.8 |
| Exposed | 773 | 10.5 | 12.1 |
| Partially Buried | 2046 | 25.0 | 20.4 |
| Buried | 407 | 36.4 | 15.0 |
| 1 protein coord | 892 | 2.5 | 4.0 |
| 2 protein coord | 1219 | 30.7 | 27.4 |
| 3 protein coord | 458 | 53.9 | 18.9 |
| BB only | 818 | 14.7 | 10.6 |
| SC only | 814 | 19.8 | 18.4 |
| BB+SC | 1070 | 42.8 | 23.3 |

<sup>1</sup>3-D RISM water positions with placevent algorithm<sup>14</sup> using RDF cutoff of 10.2 and trimmed within 6 Å of CB atoms (CA for GLY) of interface residues
<sup>2</sup>Three groups of categorization of type of predicted water molecules. First, waters are classified 'buriedness' based on number of amino acid neighbors ( $n_{C\beta}$ ) with $C\beta$ within 10 Å. Exposed: $n_{C\beta} \leq 15$ ; partially buried: $15 < n_{C\beta} \leq 25$ ; buried: $n_{C\beta} > 25$ . Second, classification by 1, 2, or 3 protein coordination partners within 3.2 Å. Finally, by type of coordinating protein atoms with 3.2 Å of the water O atom: at least two backbone only (BB only), side chain only (SC only) or a mix of backbone and side chain coordination (BB+SC).
<sup>3-4</sup>Percent and number of specific types of waters recovered using recovery criteria described in *Methods*, averaged over three runs.

### Details on Derivation and Use of Statistical Potential

A total of 14,053,883 water/protein measurements from 6,342 structures of the Top8000 database were used to determine the probability of finding water molecules at particular distances and angles from 28 different protein polar groups. Probability matrices, P(d,θ), were generated by binning distance and angle measurements with bin intervals of 0.25 Å and 7.5°, respectively, followed by normalization and smoothing via convolution with a Gaussian kernel. The final potential for each polar group takes the form of –log(P(d,θ) / P(d,θ)_ref_), where the reference is the distribution of water about a non-polar group (the β-carbon of alanine). The final potential maps, as illustrated in Fig S3, were flattened to zero beyond 4 Å and 90° to only include first solvation shell effects. Additionally, testing has shown that normalized potentials result in better recall of natives, thus all point water potential minimums were set to a value of −1.5, which roughly corresponds to the score awarded to a single hydrogen bond in the full-atom Rosetta energy function.

The initial placement of water molecules to be scored by the point water potential come from two sources. First, possible water sites about backbone polar atoms are obtained from the same statistics used to develop the point water potential. From the experimental data set, all waters within 3.1 Å of a backbone carbonyl or nitrogen group were clustered into discrete representations of the entire statistical distribution based on k-means clustering. Ultimately, positions of 77,798 water molecules about C=O groups in the data set were clustered down to 10 most-probable solvation sites (illustrated in Figs S4&5), while only a single site is used for the NH backbone group.

Given that the backbone is fixed in most Rosetta protocols, a strategy of building possible water sites based on statistics from the PDB is very efficient and accurate. However, this strategy becomes less feasible when applied to identifying potential solvation sites about the polar groups of side chains. Given the large number of rotameric states available to most polar side chains, the statistical distribution of water sites becomes very disperse. To overcome this problem, we instead implement a method in which idealized water positions about polar groups, as defined by Yanover and Bradley^23^, are accumulated for all rotamers available to each amino acid that is being considered for solvation. The resulting set of potential water sites is then reduced to only keep the average position of pairs of waters within 0.75 Å of each other that originate from two different residues, with the goal of obtaining possible water sites that could bridge the interaction between two side chains. The resulting positions are further culled by removing duplicate positions within 1.0 Å of each other. The final positions are then clustered with a 3.0 Å radius to form rotamer sets for each grouping.

Using the statistical point water potential to solvate a protein surface or interface involves a modified version of the standard Monte Carlo (MC) packing algorithm used in Rosetta. For each position on a protein to be solvated, clouds of solvation sites are built off a fixed backbone. Each collection of solvation sites is treated as a new residue composed of point water ‘rotamers’. During the packing simulation, a single rotamer is selected from the entire rotamer set including both point waters and side chains. The rotamer is applied to the pose, scored, then accepted or rejected based on the Metropolis criterion. Since a majority of the water rotamers are expected to score poorly, each water residue is assigned an extra ‘virtual’ state which is sampled 50% of the time a water rotamer is randomly selected to help convergence of the simulation.

Test simulations with the point water potential have shown that the most relevant water states were visited during low temperatures. Therefore, long simulations at a single low temperature (RT = 0.3 kcal/mol) were used as an alternative to the default simulated annealing protocol of Rosetta. Periodic temperature spikes at 100 K serve to scramble the overall conformation and allow for better convergence and reproducibility. For a packing simulation with a total number of rotamers equal to nrot, data is collected at the low temperature for 5*nrot MC steps, followed by a high temperature spike for nrot steps. Before data collection is resumed at the low temperature, a burn-in period of nrot steps is implemented. In total, data is collected for 50 cycles of temperature spiking.

During the packing simulation, the dwell time for each point water rotamer is recorded and ultimately normalize to the total number of low-temperature MC steps. Those positions with a dwell time below a specified cutoff are discarded and the remaining positions are clustered to dwell time-weighted centroid positions. A second cutoff is used to remove clusters with a cumulative dwell time below a certain threshold. The final centroid positions are then converted to full-atom water molecules. The water molecules, which sample positions in rotational space and again include a virtual state, are packed one final time to make use of the full Rosetta energy function, further discriminating between the true positive positions from the false.

### Description of Training Tasks and Results on Test Sets

Three different classes of training tasks were used in the parameterization of the *Rosetta-ICO* energy function, which were the same used for the development of the *REF2015* energy function. While the tasks are described in detail in the REF2015 paper (Park et al, 2016), they are summarized below.

Briefly, the three categories include: structure prediction, sequence design, and high-resolution structure recovery. For the structure prediction tests, two sets of monomeric protein folding energy landscapes are used to evaluate the ability of the new energy function to properly identify near-native structures from decoys. For sequence design, individual residues of protein monomers as well as residues at protein-protein and protein-ligand interfaces are mutated to all 20 amino acids, followed be re-optimization of the structure with a fixed backbone, and scoring with an entropy-weighted profile recovery metric. Additionally, correlation to mutational ΔΔG were computed as a further evaluation metric. Finally, for the high-resolution structure recovery test, atom-pair distributions are compared for structure refined with two rounds of the FastRelax protocol of Rosetta in Cartesian space using the target score functions.

For structure prediction tasks, we report the Boltzmann weight of near-native decoys and the percent of native structures recovered; for sequence design, we report the entropy-weighted native recovery except for ΔΔG which reports the Pearson correlation; and for the high-resolution structure recovery, we report the relative error. The results are presented in Table S2. For all tests except high-resolution recovery, higher values are indicative of improved performance. Overall we see similar performance between REF2015 and Rosetta-ICO on monomeric structure prediction and protein design tasks. Looking at the “percent success” metric, we see a modest improvement in decoy discrimination, from ~ 62% to 65%; in all other metrics the two score terms are comparable. Therefore, we believe this is a reasonable general-purpose energy function.

**Table S2.** Performance of Energy Functions on Training Task Test Sets.

| Tasks |  | *REF2015* | *Rosetta-ICO* |
| --- | --- | --- | --- |
| structure prediction | decoy discrimination, set 1 | 0.705 (57.1) | 0.728 (64.9) |
|  | decoy discrimination, set 2 | 0.781 (67.2) | 0.761 (65.6) |
| sequence design | protein monomer | 0.282 (47.0) | 0.290 (45.6) |
|  | protein-protein interface | 0.316 (51.0) | 0.314 (50.9) |
|  | protein-ligand interface | 0.425 (65.5) | 0.423 (59.2) |
| | mutational $\Delta\Delta G$ | 0.743 (72.9) | 0.733 (71.3) |
| High-resolution structure recovery | atom-pair distribution | 0.00796 | 0.00949 |

### High-Resolution Protein-Protein Interface Set (153 cases)

1DJ7, 1DPJ, 1E3D, 1F60, 1G8K, 1GO3, 1H32, 1JAT, 1JIW, 1L6X, 1LQV, 1MCT, 1MCV, 1NRJ, 1P57, 1PK1, 1PQ1, 1PXV, 1R0R, 1R8S, 1RDQ, 1SVD, 1T0P, 1T6G, 1WMH, 1WQJ, 1Z3E, 1ZLH, 2A5D, 2A9K, 2AQ2, 2BCG, 2BLF, 2BNU, 2D7C, 2F91, 2FCW, 2FHZ, 2FOM, 2H7Z, 2HQH, 2HQS, 2IEJ, 2NL9, 2NNU, 2NPT, 2NW2, 2OMZ, 2OZN, 2P1M, 2QME, 2QWO, 2R25, 2UYZ, 2V52, 2V9T, 2VLQ, 2VN6, 2VSM, 2VU8, 2WBW, 2WWX, 2WY3, 2WY8, 2X83, 2XFG, 2XPP, 2Y5F, 2YLE, 2Z30, 2Z7F, 2ZA4, 2ZFD, 2ZSI, 3AWU, 3BC1, 3C9A, 3CJS, 3D3B, 3DBO, 3DRA, 3DSS, 3EGV, 3F1N, 3FJU, 3FPN, 3FPU, 3G9A, 3GJ3, 3GMO, 3H7H, 3H8K, 3IXS, 3K2M, 3K9O, 3KF6, 3KNB, 3KSE, 3KTA, 3KYJ, 3L51, 3LXR, 3ML1, 3MMY, 3MXN, 3N1F, 3N4I, 3NHE, 3P73, 3P8B, 3P95, 3PRO, 3QN1, 3RNQ, 3SBT, 3SHG, 3VU9, 3VZ9, 3WHT, 3WN7, 3ZEU, 3ZKQ, 4A94, 4AG1, 4APX, 4CBU, 4CRU, 4CRW, 4DH2, 4DRI, 4EQA, 4FBJ, 4G1Q, 4G7X, 4GFT, 4GVB, 4HDR, 4HI8, 4HT3, 4IUC, 4JZZ, 4K12, 4K5A, 4KT3, 4KT6, 4KVG, 4L2I, 4LGR, 4LV5, 4M6B, 4MBG, 4N9O, 4NBX

### Protein-Ligand Docking Set (46 cases)

6376 total models including both near-natives and decoys

1GPK, 1HNN, 1JLA, 1KE5, 1KZK, 1L2S, 1M2Z, 1N1M, 1N2J, 1N46, 1NAV, 1OF1, 1OF6, 1OPK, 1OWE, 1P62, 1PMN, 1Q1G, 1Q41, 1R55, 1S19, 1S3V, 1SQN, 1T40, 1T46, 1TOW, 1TT1, 1TZ8, 1U1C, 1U4D, 1UNL, 1UOU, 1V0P, 1VCJ, 1W2G, 1X8X, 1XM6, 1XOQ, 1Y6B, 1YV3, 1YVF, 1YWR, 1Z95, 2BM2, 2BR1, 2BSM

### Protein-Protein Docking Set (53 cases)

59,738 total models including both near-natives and decoys

1A2K, 1AHW, 1AKJ, 1AVX, 1BJ1, 1BUH, 1BVK, 1DFJ, 1E6E, 1EAW, 1EZU, 1F34, 1F51, 1FFW, 1FQ1, 1FSK, 1GLA, 1GPW, 1H1V, 1HCF, 1IB1, 1J2J, 1JMO, 1JZD, 1KKL, 1M10, 1NCA, 1NSN, 1OPH, 1QA9, 1R6Q, 1RV6, 1SYX, 1US7, 1XU1, 1YVB, 1ZHI, 2A5T, 2AJF, 2AYO, 2CFH, 2H7V, 2I9B, 2JEL, 2MTA, 2O3B, 2OOB, 2PCC, 2UUY, 2VIS, 3CPH, 3D5S, 7CEI

### Astex Diverse Subset (67 cases)

1U4D, 1XOZ, 1J3J, 1Q41, 1OF6, 1S3V, 1UOU, 1T46, 1IA1, 1N2J, 1OYT, 1K3U, 1GPK, 1M2Z, 1W1P, 1SQN, 1N46, 1R9O, 1Z95, 1V4S, 1OPK, 1L7F, 2BM2, 1JLA, 1N2V, 1TOW, 1N1M, 1U1C, 1OF1, 1SJ0, 1HWI, 1W2G, 1X8X, 1TZ8, 1YVF, 1S19, 1YV3, 1VCJ, 1NAV, 1SG0, 2BSM, 2BR1, 1UNL, 1LPZ, 1Q4G, 1V48, 1Q1G, 1P62, 1IG3, 1HNN, 1TT1, 1MEH, 1G9V, 1KZK, 1KE5, 1GM8, 1PMN, 1YWR, 1V0P, 1YGC, 1T9B, 1OWE, 1SQ5, 1HVY, 1T40, 1L2S,

